# Role of membrane estrogen receptor alpha on the positive feedback of estrogens on LH secretion

**DOI:** 10.1101/2023.03.22.533736

**Authors:** Mélanie C. Faure, Rebeca Corona, Catherine de Bournonville, Françoise Lenfant, Jean-Michel Foidart, Charlotte A. Cornil

**Author notes:** Corresponding autor ORCID: 0000-0002-5536-7753. Departamento de Neurobiología Celular y Molecular, Instituto de Neurobiología, UNAM, Querétaro, México.

## Abstract

Estrogens act through nuclear and membrane-initiated signaling. Estrogen receptor alpha (ERα) is critical for reproduction, but the relative contribution of its nuclear and membrane signaling is unclear. To address this question, we used two complementary approaches: estetrol (E_4_) a natural estrogen described to act as an agonist of nuclear ERα and a mERα antagonist and the C451A-ERα mouse lacking mERα. E_4_ dose-dependently blocks ovulation in female rats, but the mechanism underlying this effect is unknown. To determine whether E_4_ acts centrally to control ovulation, we tested its effect on the positive feedback of estradiol (E_2_) on LH secretion. In ovariectomized females chronically exposed to a low dose of E_2_, estradiol benzoate (EB) alone or combined with progesterone (P) induced a LH surge and the associated increase in the number of activated kisspeptin (Kp) and gonadotropin-releasing hormone (GnRH) neurons. However, E_4_ blocked these effects of EB when provided alone, but not when combined to P. These results indicate that E_4_ blocked the induction of the positive feedback and the associated neuronal activation in the absence of P, suggesting an antagonistic effect of E_4_ on mERα as shown in peripheral tissues. In parallel, C451A-ERα females do not show a pre-ovulatory LH surge and the associated activation of Kp and GnRH neurons in response to EB unless they are treated with P. The similarity of the responses of C451A-ERα mice and wild-type females treated with E4 supports a role for membrane-initiated estrogen signaling in the EB-induced LH surge.

## INTRODUCTION

Estrogens, such as 17β-estradiol (E_2_), control ovulation at multiple levels of the hypothalamus-pituitary-gonadal axis (1). At the top of the hierarchy of events controlling ovulation is gonadotropin-releasing hormone (GnRH) produced by neurons scattered throughout the preoptic area. The pulsatile release of GnRH in the portal vein system drives the pulsatile release of gonadotropins from the anterior pituitary into the general circulation. Gonadotropins then act on the gonads to govern ovarian steroidogenesis and folliculogenesis. During most of the reproductive cycle, estrogens exert a negative feedback on gonadotropin secretion notably by their suppressing action on the activity of GnRH neurons. At mid-cycle, estrogens switch from negative to positive feedback to generate a continuous surge of GnRH and subsequently a luteinizing hormone (LH) surge which is necessary to trigger ovulation (2–4). How estrogens contribute to the initiation of the preovulatory LH surge remains however unclear.

Studies based on complete estrogen receptor (ER) knock out mice identified the nuclear estrogen receptor alpha (nERα) as the primary ER involved in the central control of reproduction (5–9). Upon activation, ERα modulates the transcription of target genes either through a direct interaction with an estrogen response element (ERE; classical genomic action) on the DNA or via protein-protein interaction with another transcriptional factor (tethered genomic action, (10)). Besides these genomic actions, ERα also activate membrane-initiated signaling cascades following palmitoylation (11–14). Nuclear- and membrane-initiated responses usually follow different temporal patterns: nuclear actions generally lead to relatively slow and long lasting effects, while membrane-initiated actions typically occur within seconds to minutes, although they can also indirectly lead to transcriptional changes (15–17).

GnRH neurons do not express ERα (18). The estrogen positive feedback is mediated by ERα-expressing afferents to GnRH neurons mainly originating from the anteroventral periventricular nucleus (AVPv) (7,19). In particular, kisspeptin (Kp) neurons exert a pivotal role in translating changes in circulating estrogen concentration into changes in the activity of GnRH neurons and the generation of LH surge (20–22). Although other neuronal populations likely contribute to the regulation of GnRH neurons by estrogens, the current view posits the Kp neurons located in the AVPv as key elements of the core surge generator (23).

Whether estrogen’s regulation of LH surge involves nuclear- or membrane-initiated signaling, or a combination of both is currently unclear. Early evidence indicated that a prolonged exposure to high circulating estrogen concentrations is required to elicit a LH surge (24,25) suggesting that classical estrogen signaling is involved. This is supported by studies using a knock in approach to selectively restore ERE-independent signaling in ERαKO mice which showed that ERα ERE-independent activity alone is not sufficient to restore E_2_-induced changes in the firing rate of GnRH neurons or the LH surge (26,27). However, that transcriptional signaling is required does not preclude a role of membrane-initiated signaling. Moreover, evidence exists indicating that GnRH neurons are influenced by membrane-initiated estrogen signaling (Reviewed in (28–30)). Some studies have shown direct effects of E_2_ on GnRH neurons that depend on the activation of ERβ (31–33) or another receptor, such as the STX receptor (34) or GPER1 (33). ERα-mediated indirect actions have also been reported (32,35,36). Specifically, E_2_ acutely modulates GABAergic input to GnRH neurons leading to rapid changes in their activity (32,36). Interestingly, subpopulations of GnRH neurons were identified that display activated or inhibited firing in response to E_2_ (35). Membrane ERα was also shown to stimulate the activity of Kp neurons and contribute to the regulation of Kp expression in the AVPv (37). Yet the impact of these effects of membrane-initiated action of ERα on preovulatory LH surge remains unclear.

The present study took advantage of two complementary approaches. First, the C451A-ERα mouse, a knock in mouse model with a selective loss of function of membrane signaling of ERα due to the point mutation of palmitoylation site Cys451 into an alanine. This mouse model provides a powerful tool to dissociate the two modes of action of estrogens, i.e. action on either mERα or nERα (38). C451A-ERα females were described as infertile based on the absence of litter in the nest despite mating for several months and show abnormal ovaries with hemorrhagic cysts and no corpus luteum indicative of anovulation (38). Similar observations were reported in the nuclear-only ERα (NOER) mice in which vaginal smears did not show the cornified epithelial cells typically observed during estrus (Pedram et al., 2014). Although this infertility largely relies on placental defects during development (39), this model offers the opportunity to assess the role of membrane-initiated action of ERα on the positive feedback of estrogens. Second, estetrol (E_4_) is an estrogen exclusively synthesized in human fetal liver which selectively binds ERs (ERα and ERβ) but with a lower affinity than E_2_ (40). Interestingly, E_4_ induces estrogenic actions via the activation of the nuclear ERα signaling pathway (for example in uterus (41), vagina (42), cardiovascular system (43)), while it has been described to antagonize membrane ERα in different tissues, including in the endothelium and in breast cancer cells ((41,44)). E_4_ inhibits ovulation when administered alone in rats (45) and when combined with progesterone in humans (46,47). Combined with drospirenone, E_4_ is now included in an oral contraceptive formulation (48). Yet its mechanism of action on the hypothalamic-pituitary-gonadal (HPG) axis remains unknown.

## MATERIALS AND METHODS

### Animals and general procedures

All wild-type (WT-ERα) and C451A-ERα mice of the CD1 strain, obtained by backcrossing the original C451A-ERα mice (C57Bl/6) into the CD1 background (38), were housed and bred in the animal facility of the University of Liège (Animalerie Centrale, agreement number LA1610002). Mice were genotyped by PCR analysis of DNA collected from the tail as described previously (38). Mice were weaned at around 3 to 4 weeks of age and housed in same-sex cages. All animals had free access to food and water. The room temperature was maintained at 24±2°C. Animals were housed under a reversed 12h light/dark cycle (lights on at 0100h) when tested for positive feedback (Exp. 1 & 2), or (lights off at 0800h) when tested for estrous cycle & sexual behavior (Exp. 4 & 5) and under normal 12h light/dark cycle (lights on at 0700h) when tested for spontaneous ovulation and gestation success (Exp. 3). All experimental procedures were in accordance with Belgian laws on the “Protection and Welfare of animals” and on the “Protection of experimental animals” and were approved by the Ethics Committee for the Use of Animals at the University of Liège (Protocols #1620 and #1340).

### General procedures

#### Surgery

Between 2 and 3 months of age, females were bilaterally ovariectomized (OVX) under general anesthesia using a mixture of Domitor (Domitor, Pfizer, 1 mg/kg) and medetomidine (Ketamine, 80 mg/kg) administered subcutaneously (s.c.). In some experiments, animals were implanted at the time of ovariectomy with a subcutaneous Silastic® capsule filled with E_2_. At the end of surgery, medetomidine-induced effects were antagonized by atipamezole (s.c., Antisedan, Pfizer, 4 mg/kg) to accelerate recovery.

#### Hormones

17β-Estradiol (E_2_, E8875), β-Estradiol-3-Benzoate (EB, E8515) and Progesterone (P, P0130) were purchased from Sigma-Aldrich and dissolved in sesame oil, used as vehicle, unless stated otherwise. EB (s.c., 1µg) and P (s.c., 500µg) were injected s.c. (49), while E_2_ (1µg diluted in 7.35 µl of sesame oil/20g of body weight) was provided through subcutaneous Silastic® capsules (inner diameter: 1.02mm; outer diameter: 2.16mm; Dow Corning) (50). Estetrol (E_4_) was provided by Mithra Pharmaceuticals and dissolved in sesame oil without ethanol (s.c., 0.2mg) or dissolved in sesame oil with 5% ethanol at 6µg, 20µg, or 60µg (s.c.) (Exp. 3 to 5)) as E_2_ (0.6µg (Exp. 3) or 0.5µg (Exp. 4 and 5)). Unless stated otherwise, treatments were counterbalanced across housing cages such that each/every cage contained animals with different treatments.

#### Blood collection

Depending on the question and the method used for blood analysis, blood drops, blood sample or trunk blood were collected.

For repeated sampling of blood drops on a same day or assay with ultra-sensitive immune-enzyme assays (EIA; Exp. 1-part1 (p1)), blood was collected using the repetitive tail-tip blood sampling described previously (51). Briefly, mice were habituated to handling for a few minutes while massaging the tail every day during 2 or 3 weeks (51). For blood drop collection, a single excision of the tail tip was made with a razor blade. When females were OVX (regardless of whether they were treated with EB and/or P), one blood sample (5.2µl) was collected with a pipette and immediately diluted in 98.8µl phosphate buffered saline with 0.05% of tween20 (PBST), quickly frozen in dry ice and stored at −80°C until further use. In Exp. 1-p1, blood drop samples were collected every 30 min for 4 hours.

For Exp. 1-p2, mice were placed under a red-lamp to increase the dilatation of blood vessels and were restrained in the immobilizing cage where a single excision of the tail with a razor blade was made. Blood (200µl) was collected in heparinized microhematocrit capillary tubes filled by capillarity. The tail was massaged to facilitate blood dripping. Blood was stored in a 1.5ml microfuge tube containing a drop of heparin (Leo, 012866-08, 5000 U.E/ml). Blood was centrifuged 10 min at 1500g at 4°C, the plasma was collected and stored at −80°C until quantification by radioimmunoassay (RIA).

At the end of experiments (Exp. 1-p2 and Exp. 2), trunk blood was also collected in 1.5ml microfuge tube containing a drop of heparin. Plasma was collected as previously and stored at −80°C until further use.

#### LH assay

Two methods were used to assay LH: an ultra-sensitive sandwich ELISA (52) and a classical RIA (53). The ultra-sensitive sandwich ELISA was used for blood drops (Exp. 1-p1 and p2 (day 38)) while the RIA was used for all the other type of blood samples (Exp. 1-p2 and Exp. 2).

We used the sensitive sandwich ELISA previously described and validated (Steyn et al 2013) with few modifications (54). Briefly, 96-well high-affinity binding microplates (9018, Corning) were coated with 50µl of a monoclonal antibody directed against bovine LH beta subunit (1:1000; 518B7; RRID: AB_2665514, University of California, UC Davis) and incubated overnight at 4°C. Unspecific binding was blocked by incubating each well with 200µl of blocking buffer for 24h at 4°C. Samples (50µl) and LH standards (50µl; generated by serial 2 fold dilution of mouse LH starting at 400pg/well until 0,19pg/well, AFP-5306A, National Institute of Diabetes and Digestive and Kidney Diseases – National Hormone and Pituitary Program [NIDDK-NHPP]) were incubated for 2h before adding 50µl of detection antibody (1:10000; polyclonal antibody, rabbit LH antiserum, AFP240580Rb; RRID:AB_2665533, NIDDK-NHPP) for 1.5 hours at room temperature (RT). A horseradish peroxidase-conjugated polyclonal Goat Anti-Rabbit antibody (50µl, 1:2000; P0448, Dako; RRID:AB_2617138) was added in each well for 1.5 hours at RT. Then, the substrate of the peroxidase (100µL, 3,3’,5,5’-tetramentylbenzidine solution; 1-StepTM Ultra TMB-ELISA, 34029, Thermo Scientific) was added in each well for 10 to 25 minutes at RT and in darkness. The reaction was stopped by 3M HCl (50µl). The absorbance of each well was read at a wavelength of 450nm and at a wavelength of 650nm (background). The optical density (OD) obtained at 650nm was substracted from this obtained at 450nm. The amount of LH present in each well was determined by interpolating the resulting OD of unknown samples against a non-linear regression of the OD of the LH standard curve (GraphPad Prism 8). Standards were run in duplicate and yielded a non-linear curve fitting with a R^2^ >0.95. The sensitivity of the assay was 0,03ng/ml. All samples from a same mouse were assayed on the same plate and genotypes and treatments were counterbalanced within plates. The intra- and inter-assay coefficients of variation were less than 10% and 15%, respectively.

The RIA consisted of a double antibody method with reagents provided by the National Institutes of Health (Dr. A. F. Parlow, National Institute of Diabetes and Digestiveand Kidney Diseases (NIDDK), National Hormone and Peptide Program, Torrance, CA) (50). LH was detected by a rat LH-I-10 (AFP-11536B) labeled with ^125^I and precipitated with a Rabbit anti-mouse LH (AFP-240580; RRID: AB_2784499). Mouse LH reference preparation (AFP-5306A) was used to prepare the standard curve. The intra- and inter-assay coefficients were less than 10 and 7% respectively and the sensitivity of the method was set at 4 pg/100 ml based on the lowest detectable point of the standard curve.

#### Estrous cycle

**Vaginal smears** were carried out daily between 0930 and 1030h, i.e. 1h30 to 2h30 after lights off (Exp. 4). Samples were placed to dry at 37°C on a glass slide which was then immerged in cresyl violet (5g/L) for 1min and rinsed tree times in distilled water before observation without drying. Smears were examined under a light microscope for the presence of different cell types indicative of the different stages of the cycle of the mouse (55).

#### Behavioral tests

Lordosis behavior was assessed during the dark phase of the photoperiodic cycle. Sexually experienced ovariectomized females (Exp. 5) were placed in the presence of a gonadally intact male stimulus in a Plexiglas chamber (40 cm long × 20 cm wide × 25 cm high). The responses of the female to male mounts were recorded until she had received 10 mounts or after 10 min had elapsed. The lordosis quotient was calculated by dividing the number of lordosis postures performed by the number of mounts received and multiplying this value by 100 to obtain a percentage of lordosis behavior.

Sexual partner preference was tested in a three-chamber arena that consists of a box (60 cm long × 30 cm high × 30 cm wide) divided into three equally sized compartments with open doors between each compartment (Exp. 5). The back part of both lateral compartments contained a stimulus animal that was separated from the main area by a transparent perforated partition. An intact male and a hormonally primed estrous female were each placed in one of the lateral compartments. The experimental female was introduced into the middle compartment and was observed for 10 min. The time spent by the subject in each compartment was recorded. Since no habituation to the box was performed, we did not consider the first two minutes in the final analyses. A preference score was calculated by dividing the time spent near the male minus the time spent near the female by the total time spent in both compartments. A positive score indicates a mate preference directed toward the male stimulus, whereas a negative score indicates a mate preference directed toward the stimulus female.

#### Euthanasia

Apart for Exp. 3, Exp. 4 and 5 wherein animals were killed by cervical dislocation prior to dissection of the genital tract to analyze cumulus oophori, animals were anesthetized with isoflurane and were decapitated 30 min after lights off (Exp. 1 and 2). Their brain was then removed from the skull and immersed in a solution of 0.5% of acroleine in 0.01M PBS for 2 hours at RT. For this type of fixation, brains were rinsed twice for 30 min in PBS before being transferred in 30% sucrose overnight. Brains were then frozen on dry ice and stored at −80°C until further use. All brains were cryosectioned in four series of 30µm thick coronal slices from the corpus callosum level to the end of the hypothalamus. Sections were stored in antifreeze solution and kept at −20°C.

#### Histology and Immunostaining

Brains were double labeled for Fos and Kp or GnRH. Briefly, brain sections were first rinsed three times for 5 min in 0.05M tris-buffered saline (TBS; pH 7.6) at RT. Unless mentioned otherwise, all following incubations were carried out at RT and followed by similar rinses. Sections were first incubated in 0.1% sodium borohydrate for 15 min. They were then incubated in hydrogen peroxide (H_2_O_2_, 1% for 20 min) to block endogenous peroxidase activity. Sections were blocked and permeabilized for 1 hour in normal goat serum (NGS) in TBS with 0.1% Triton X-100 (TBST) and immediately incubated at 4°C in the primary antibody against the N-terminus of human Fos (overnight, 1:2000; Rabbit polyclonal, ABE457, Millipore; RRID: AB_2631318 (56) (Exp. 1-p2); overnight, 1:2000; monoclonal antibody, sc-166940, Santa Cruz Biotechnology; RRID: AB_10609634 (Exp. 2)) in NGS and TBST. Sections were then incubated for 2 hours in a goat anti-rabbit biotinylated antibody (111-065-003; RRID: AB_2337959; Jackson) followed by one hour in the AB complex solution (PK-6100 Vector Laboratories) diluted at 1:400 or 1:800 (for Fos when followed by GnRH labeling or for Fos when followed by Kp labeling, respectively). The immunoproduct was visualized with 0.05% diaminobenzidine with 0.012% H_2_O_2_ in TBS.

When double labeling, the first visualization was followed by a blockade of avidins and biotins using avidin-biotin blocking Kit (SP-2001; Vector Laboratories) for 15 min prior to an additional blocking and permeabilization step. Sections were immediately incubated overnight in a polyclonal rabbit antibody directed against GnRH-I (1:400, polyclonal, #20075, Immunostar; RRID: AB_572248 (57)) or twice overnight in a rabbit antibody directed against mouse Kp (1:10000; rabbit polyclonal, Ac566 kindly provided by Isabelle Franceschini and Massimiliano Beltramo, INRA, Nouzilly, Tours, France; RRID: AB_2296529 (58)) in NGS and TBST. Sections were then incubated in a goat anti-rabbit biotinylated secondary antibody (111-065-003; Jackson Immunoresearch). Finally, the immunoproduct was visualized by a last incubation in the substrate of the Vector SG Peroxidase Substrate Kit (SK-4700; Vector laboratories). After final rinses, sections were mounted on microscope slides and coverslipped with Eukitt (Sigma-Aldrich, Bornem, Belgium).

#### Image analysis

**Brains** - The number of single labeled Kp-immunoreactive (ir) neurons or the number of Kp-ir and GnRH-ir neurons co-labeled with Fos was analyzed by direct observation at 40x magnification using Leica DMRB microscope. The number of Kp-ir cell bodies was investigated bilaterally in ten consecutive brain sections (each separated by a distance of 90µm) encompassing the anteroventral periventricular nucleus (AVPv) and the rostral periventricular nucleus (PeN) continuum (corresponding to plates 29 to 35 of the Paxinos Mouse Atlas (59)). The number of GnRH-ir cell bodies was analyzed bilaterally in ten consecutive brain sections (each separated by a distance of 90µm) corresponding to plates 21 to 31 of the Paxinos Mouse Atlas (59). Kp and GnRH immunolabeling is cytoplasmic, while Fos immunolabeling is detectable only in the nucleus. All Kp or GnRH neurons detected in this region were counted and analyzed for the presence of nuclear immunostaining for Fos. The values obtained for each side of the ten sections were summed to provide a total number of Kp or GnRH expressing neurons and the percentage of Kp or GnRH neurons co-expressing the protein Fos.

### Experimental designs

#### Experiment 1 – Positive feedback

The role of mERα in the induction of LH surge was repeatedly assessed in 2 cohorts of 2 month old WT-ERα (Cohort 1: n=24; Cohort 2, n=17) and C451A-ERα (Cohort 1: n=18; Cohort 2, n=19) females. The two cohorts were subjected to the exact same protocol except that females from the second cohort were housed based on their treatment. In each cohort, females were tested twice following a paradigm of LH surge induction, i.e. by implantation of a subcutaneous capsule delivering low levels of E_2_ mimicking diestrus levels and administration of EB 7-8 days after OVX (Fig. 1A) (60,61). The first test was designed to examine the time-response profile of the EB-induced LH surge following blood sampling every 30 min for 4 hours (Part 1, day 0 to 8; (51)), while the second investigated the central activation of the circuit underlying the LH surge, namely Kp and GnRH neurons (Part 2, day 30 to 39; (49,50)).

**Fig. 1.**
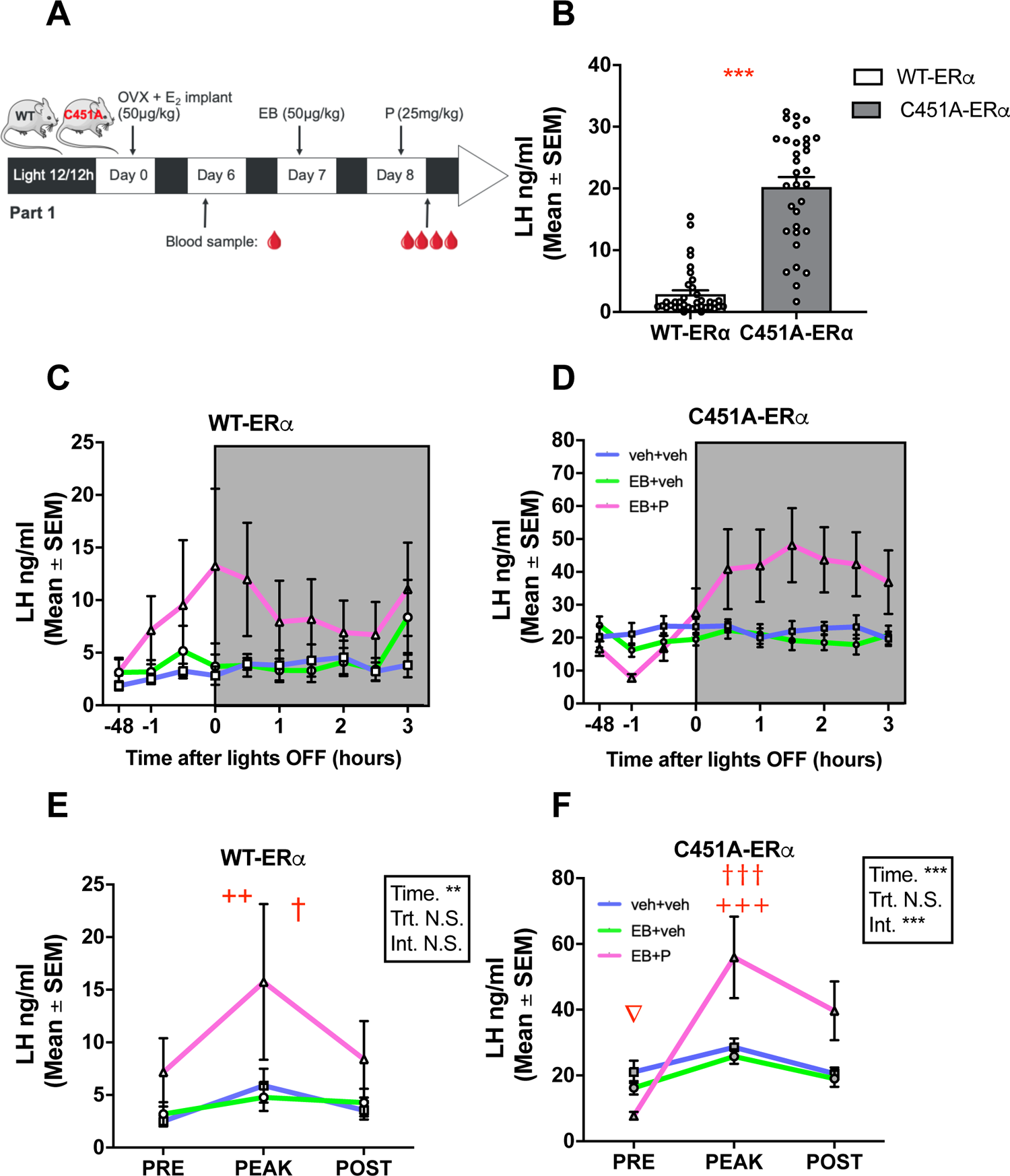
Profiles of LH changes induced by estradiol benzoate (EB) alone or in combination with Progesterone (P) in ovariectomized WT-ERα (white) or C451A-ERα (gray mice). A. Protocol used to induce a positive feedback: Females were ovariectomized (OVX), chronically treated with estradiol (E_2_) from Day 0 to day 8, injected with estradiol benzoate (EB) on day 7, and Progesterone (P) or its vehicle (Sesame oil) on day 8. B. On day 6, C451A-ERα females showed higher baseline LH levels than WT-ERα females (U=47, p<0.0001). C and D Profiles of LH levels measured every 30 min starting 1h before lights off following treatment on day 8 in WT-ERα and C451A-ERα females respectively. C. LH profiles obtained in WT-ERα mice (OVX+E_2_+veh+veh n=12, OVX+E_2_+EB+veh n=11, OVX+E_2_+EB+P n=14). D. LH profiles obtained in C451A-ERα mice (OVX+E_2_+veh+Veh n=10, OVX+E_2_+EB+veh n=11 OVX+E_2_+EB+P n=12). E. WT-ERα females showed an increased LH concentration at one time point (peak) between 0 and 2.5 hour after lights off compared to prior (day 6, pre) and during 3h after lights off (post) regardless of treatment (2-way ANOVA; Tukey post hoc test, following significant time effect: +, ++, +++, p<0.05, 0.01, 0.001 time effect compared to pre time; †, ††, †††, p<0.05, 0.01, 0.001 time effect compared to post time). F. Only EB+P induced an increased LH concentration in C451A-ERα females at one time point (peak) between 0 and 2.5 hour after lights off compared to prior (day 6, pre) and during 3h after lights off (post) (2way-ANOVA; Tukey post hoc test, following significant interaction: +++, p=0.001 compared to pre time same treatment; †††, p=0.001 compared to post time same treatment; ∇ p<0.05 EB+P pre time condition compared to post time same treatment;. Symbols in the statistical boxes: *, **, ***, p<0.05, 0.01, 0.001; N.S., non-significant).

Briefly, females were OVX and implanted with a subcutaneous capsule containing E_2_ (1µg). A first blood sample was collected on day 6 (D6) post-ovariectomy between 0820h and 0900h (3 µl immediately diluted in 57 µl of PBST for EIA). Females of each genotype were subdivided in three groups of equal size subjected to 3 different hormonal treatments s.c.: veh+veh, EB+veh and EB+P. On day 7 (1000h), they were injected with EB or its vehicle (veh). On day 8 (1000h), they were injected with P or veh 3 hours before lights off, while females that had received veh on day 6 received veh again. Blood sampling was then carried out every 30 min for 4 hours starting 60 min before lights off. All samples were assayed in duplicate. Three to 7 days later, their implant was removed, and they were treated every 3 or 4 days with EB until the beginning of the second part.

Part 2 started 30 days after part 1. Females were re-implanted with a new subcutaneous E_2_ implant. Two blood samples were collected on day 38 between 0800h and 0900h: 5.2 µl immediately diluted in PBST for EIA and 200µl for plasma collection and RIA. Females were then treated with veh or EB at about 1000h. The next day (day 39), veh or P was injected 3 hours before lights off. Mice were anaesthetized with isoflurane 30 min after lights went off and euthanized by rapid decapitation. Trunk blood was collected, extracted for plasma as described above, and assayed by EIA. Brains were fixed in 0.5% acrolein (Fig. 2A).

**Fig. 2.**
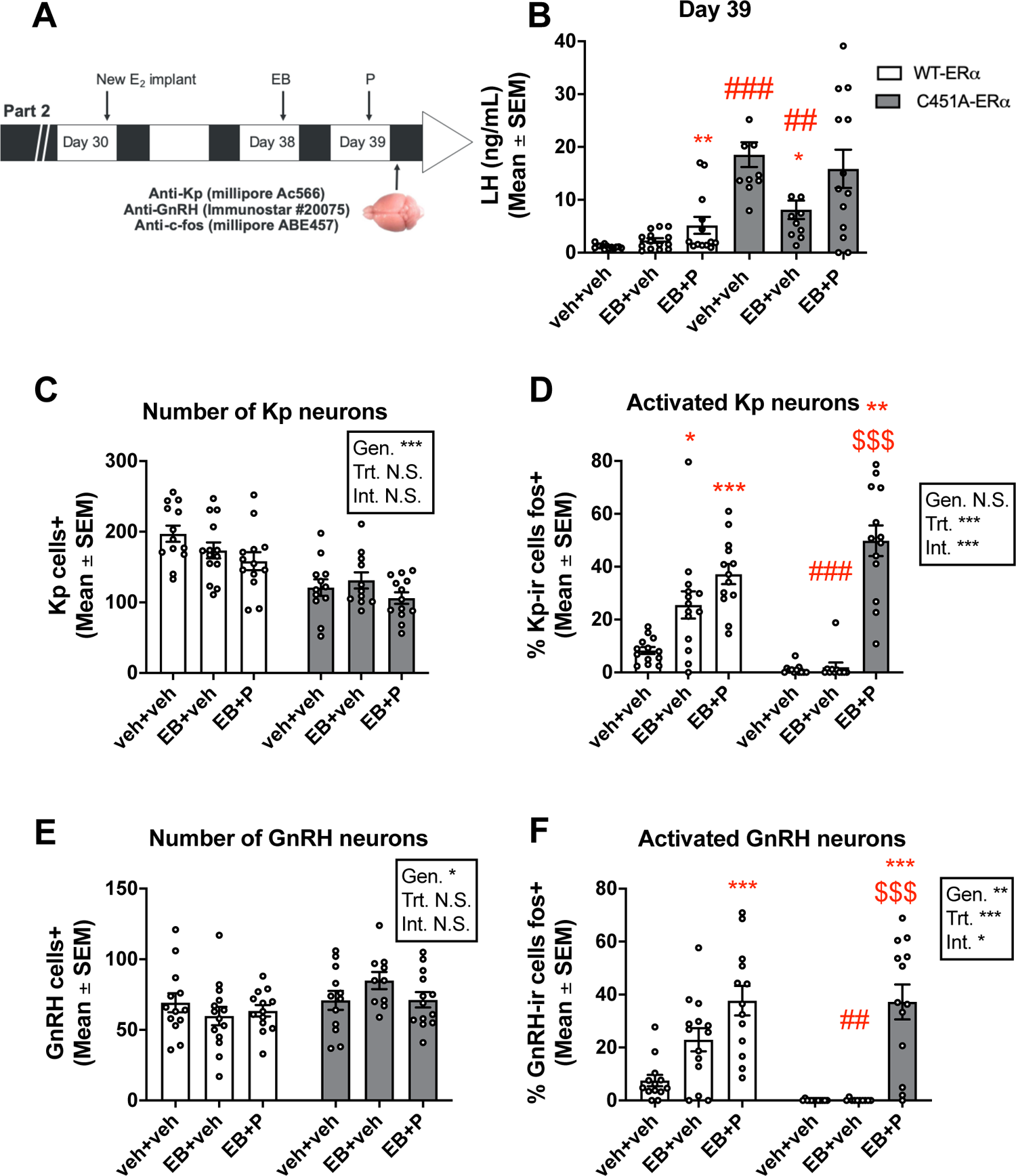
Effect of mERα absence on the positive feedback of estrogens on LH concentration and the activation of the associated neurocircuits. A. Protocol used to induce positive feedback: following a first round of injections to induce the positive feedback (see *figure 1*), the E_2_ implant were replaced by a new one on day 30, and females were treated again with veh+veh, EB+veh or EB+P on day 38 and 39. Blood and brains, were collected 30-60 minutes after light off. B. In WT-ERα females (white), EB+P, but not EB+veh, induced a significant rise in LH (Kruskal Wallis test: **p<0.01 vs veh+veh) while in C451A-ERα females (gray), EB+veh induced a significant reduction in LH (Kruskal Wallis test: *p<0.05 vs veh+veh; Mann Whitney tests: ##, ### < 0.01, 0.001 vs WT-ERα same treatment). C and E. Number of Kisspeptin (Kp) neurons counted in RP3V (AVPv + PeN;C) and GnRH neurons counted in POA (E, Two-way ANOVA) for each treatment and genotypes. D and F. Percentage of Kp and GnRH neurons co-expressing Fos in RP3V and POA respectively. D. Kp neurons are activated by both EB and EB+P in WT-ERα, while only EB+P induced such activation in C451A-ERα (Two-way ANOVA; * and ***, p<0.05 and 0.001 vs veh+veh same genotype; $$$, p <0.0001 vs EB+veh same genotype; ###, p<0.001 vs same treatment in WT-ERα). F. GnRH neurons are activated by both EB and EB+P in WT-ERα, while only EB+P induced such activation in C451A-ERα (Two-way ANOVA; ***, p<0.001 vs veh+veh same genotype; $$$, p <0.0001 vs EB+veh same genotype; ##, p<0.01 vs same treatment in WT-ERα. Symbols in the statistical boxes: *, **, ***, p<0.05, 0.01, 0.001; N.S., non-significant.

#### Experiment 2 – E_4_ and Positive feedback

This experiment investigated the effect of E_4_ on the induction of LH surge in WT-ERα females (n=45) subjected to a classical paradigm of induction of the LH surge by administration of EB with or without P in OVX females chronically exposed to low estrogen levels mimicking diestrus levels (Fig. 3A) (60,61). Briefly, females were OVX and implanted with a subcutaneous E_2_ capsule. Prior to treatment, one blood sample (200µl) was collected on day 8 after ovariectomy. Females were subdivided in 5 groups and subjected to 5 different hormonal treatments: veh+P, EB+veh, EB+P, EB+E_4_ and EB+E_4_+P. On day 8 (1000h), they were injected with veh, EB or EB+E_4_. On day 9 (1000h), they were injected with P or veh 4 hours before lights off. Females were euthanized by rapid decapitation within 1 hour after lights off, trunk blood was collected and the brain was dissected out of the skull and fixed in 0.5% acrolein (49,50).

**Fig. 3.**
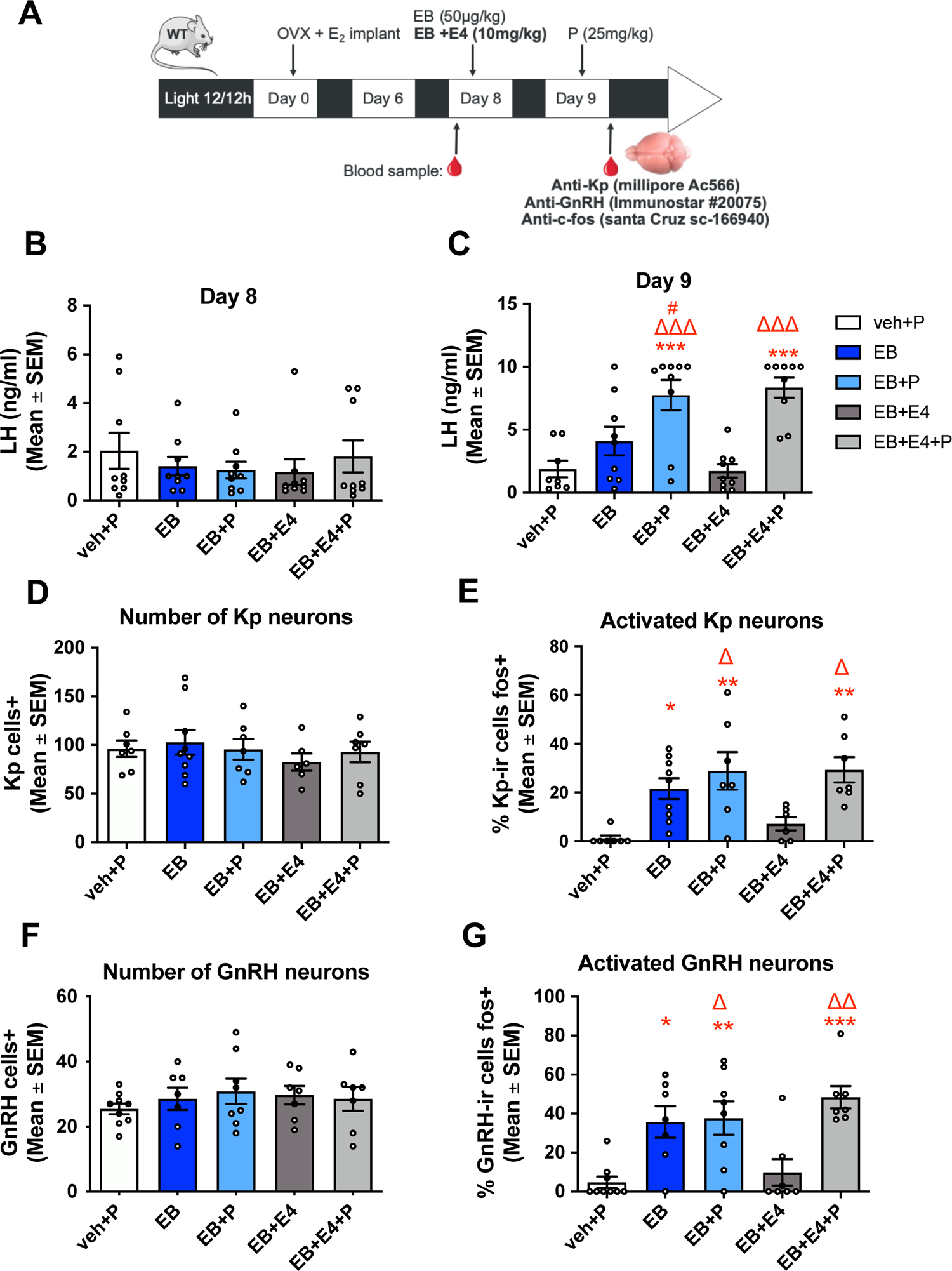
Effect of estetrol on the LH surge induced by estradiol the neuro-circuits underlying this response. **A.** Protocol used to induce a positive feedback. WT mice were ovariectomized (OVX) on day 0, treated with subcutaneous estradiol implant (E_2_) from day 0 to day 9, and injected on day 8 with estradiol benzoate (EB) alone or combined with estetrol (E_4_) or their vehicle (Sesame oil) and on day 9 with progesterone or its vehicle (Sesame oil). Blood samples were collected prior to treatment administration on day 8 and within 1 hour of lights off on day 9, when brains were also collected for immunohistological analyses. **B.** LH levels did not differ between groups on day 8. **C.** Females treated with EB alone or EB+E_4_ did not show a LH surge unless they were treated with P (EB+P and EB+E_4_+P). **D and F.** The number of Kisspeptin (Kp) neurons in RP3V (AVPv + PeN, **D**) or GnRH neurons in POA (**F**) did not differ across treatments. **E and G.** The percentages of Kp (**E**) and GnRH (**G**) neurons co-expressing Fos were higher in females treated with EB, EB+P and EB+E_4_+P than with veh+P and EB+E_4_. All data were analyzed by one-way ANOVA followed by a Tukey’s post hoc test when significant: *,**, and *** p<0.05, 0.01 and 0.001 vs veh+P; #, p<0.05 vs EB; Δ, ΔΔ and ΔΔΔ, p<0.05, 0.01 and 0.001 vs EB+E_4_.

#### Experiment 3 – Effect of E_4_ on spontaneous ovulation and gestation success

This experiment investigated the effect of E_4_ on the capacity of females to become pregnant and carry a pregnancy to term as well as the number of pups born. Gonadally intact 3 month old females (n=53) were divided in four groups which were injected twice daily with E_4_ (6, 20, 60µg) or veh for 5 days. Females were housed according to treatment. Three days after treatment onset, females were placed overnight with a male (2 females per male), checked for a vaginal plug on the next morning (day 4, 0830h). The percentage of plugged females, the number of pregnant female (based on the number of females that gave birth, detectable parturition with pups alive or blood in the nest) and the number of pups per nest were counted. The same experimental design was run three times with an interval of 2 weeks between runs. Importantly, each female received a different treatment in each run resulting in a sample size of 36 in each of these four conditions. A fifth group of females treated with E_2_ (0.6µg; n=16) was however added in the third run as a positive control.

To determine whether treatments alter ovulation, the same females (then aged 7 months) were (n=52) used again and divided in three groups which were injected twice a day with E_4_ (60µg; n = 18), E_2_ (0.6µg; n = 17) or veh (n = 17) for three days. They were then mated overnight with a sexually experienced male once a week for 2 weeks until they showed a vaginal plug on the next day (0830h). Females with a plug were euthanized within 3 hours of vaginal investigation. The infundibulum was opened with a needle at the level of the ampula and the cumulus oophori ovulated by each horn were counted. The sum of oocyte laid by the two horns was then calculated and averaged for each treatment.

#### Experiment 4 – Effect of E_4_ on cyclicity

This experiment investigated the effect of E_4_ on the female estrous cyclicity. Gonadally intact wild-type females (n=10; 3 months old) were divided in three groups which were injected twice daily with E_4_ (60µg) or E_2_ (0.5µg) or veh for 3 days. Estrous cycles were assessed by vaginal smears performed every morning between 0930h and 1030h. Vaginal smears were analyzed on the 3^rd^ day which corresponds to the day when ovulation and sexual behavior was assessed in Exp. 3 and 5. The animals were divided into three subgroups which were repeatedly subjected to the three treatments in a counterbalanced order over a period of 3 weeks.

#### Experiment 5 – Effect of E_4_ on female sexual behavior

This experiment investigated the effect of E_4_ alone or in combination with E_2_ on two aspects of female sexual behavior, lordosis behavior and partner preference (PPT) in OVX females. Stimulus animals were prepared and include both males and females: stimuli males were kept gonadally intact while stimuli females were gonadectomized and hormonally primed before the behavioral tests (EB) 48h and 24h before the behavioral test and progesterone 3-4h before the behavioral tests). OVX females (n=24, 3 month old) primed with EB and P were given sexual experience by placing them in the presence of an intact male stimulus in sets of training sessions. First, two sessions were carried out in for 2 hours in the female’s homecage and sexual activity was evaluated by checking for the presence of a vaginal plug. For the two additional sessions the interaction occurred in a test arena (40 cm long × 20 cm wide × 25 cm high) where the number of lordosis postures displayed by in response to male mounts were recorded until she had received 10 mounts or after 10 min had elapsed. The average lordosis quotient of the two scored sessions was 78 and 68 respectively.

Females were then repeatedly tested to compare the effect E_2_ (s.c., 0.5µg) and E_4_ (s.c., 60µg) given alone or in combination on lordosis behavior and partner preference. Estrogens or their vehicle were injected twice a day for 2 days and P injected on the 3^rd^ day 4 hours before test. Females were tested on consecutive weeks for the different treatments, which were given in different order to three different subgroups to control for potential carry over effects.

### Statistical analysis

All statistical analyses were performed using Prism 8 (Version 8.0.0, GraphPad Software). Continuous data were analyzed by parametric unpaired student t tests and 2-way ANOVAs or by non-parametric Mann-Whitney and Kruskall Wallis tests when the normality and homoscedasticity assumptions were violated. Significant parametric and non-parametric ANOVAs were followed by Tukey and Dunn’s post hoc tests, respectively. Non-directional one-sample two-tailed t tests were also performed as an extra analysis for partner preference data in order to assess a significant preference for a side/stimulus compared to no preference (i.e. preference score of 0). Contingency data were analyzed by Fisher’s exact tests. A Bonferroni correction was applied when multiple Mann-Whitney tests and Fisher’s exact tests were applied to a data set and the resulting p value is called adjusted p value (p_adj_). Due to technical issues such as the loss or the degradation of sections during processing, the final sample size may differ from the initial number of samples collected, thus explaining the variability in the degrees of freedom between analyses of samples originating from the same experiments. Results were considered significant when p<0.05. All results are represented as means ± SEM unless mentioned otherwise.

## Results

### Are C451A-ER*α* mice able to show a LH surge in response to EB and is progesterone P necessary?

Although the paradigm of rising E_2_ levels can induce an LH surge in the absence of P, the addition of P yields changes of higher amplitude (62,63). Therefore, the first experiment investigated the role of mERα on the LH surge profile induced by EB combined or not with P. OVX females were implanted with a capsule delivering low E_2_ amounts mimicking circulating E_2_ levels at diestrus (60) and blood was collected by tail tip blood sampling every 30 min for 4 hours starting 1 hour prior to lights off (Exp.1). This experiment was conducted in two cohorts of mice, subjected to the exact same protocol, whose data were pooled. First, looking at baseline LH levels (6 days after OVX and implantation of a subcutaneous capsule delivering low levels of E_2_), C451A-ERα females showed significantly higher LH levels than their WT-ERα littermates (C451A-ERα, Median = 21.63, n= 36; WT-ERα, median= 1.451, n= 42; U=47, p<0.0001 Fig 1B), indicating that C451A-ERα females may present some impairment of the negative feedback.

The qualitative analysis of the average profiles of LH concentration measured every 30 min on day 8 indicated that C451A-ERα females exhibited much higher circulating LH than their WT-ERα counterparts. Despite this major difference, EB+P resulted in an increased LH concentration, while no surge was induced in the control condition (veh+veh) in both WT-ERα and C451A-ERα mice (Fig 1C-D). However, no increase in LH levels was observed following EB alone, regardless of the genotype. In WT-ERα, LH began to rise before lights off, peaked at lights off and slightly decreased afterwards while remaining elevated for the next 3 hours (Fig 1C). In C451A-ERα mice, the LH surge began with a slight delay compared to WT-ERα, peaked 30 min after lights off and remained elevated for the next 2.5 hours (Fig 1D). Interestingly, in C451A-ERα females treated with EB+P and EB+veh, the first time point (−1 hour) shows a clear decrease in LH concentration compared to the measure taken 48 hours earlier (Day 6) potentially reflecting a negative feedback exerted by EB.

For analysis purposes, the highest LH concentrations obtained between 0h and 2.5h after lights off (Peak) in each animal were averaged across females and compared to the concentration measured 48h before (Pre) and 3h after (Post) lights off (Fig. 1E-F). Confirming the qualitative observations, no LH surge was observed following treatment with veh or EB alone in both genotypes. In WT-ERα, the analysis revealed no effect of treatment (F_2,33_ = 1.488; p=0.2405), but a time effect (F_2,66_ = 6.747; p =0.0022; Fig 1E) which results from a higher LH level measured at the peak compared to the pre (p=0.022) and post conditions (p=0.026). Despite the marked increase in LH exhibited by females treated with EB+P, there was no interaction (F_4, 66_ = 1.797; p=0.1400). In C451A-ERα, the analysis revealed no effect of treatment (F_2, 30_ = 2.087; p=0.1417), but a time effect (F_2, 60_ = 19.04; p<0.0001; Fig 1F) and an interaction between the two factors (F_4,60_ = 8.519; p<0.0001). These effects are explained by significant differences between all time points in EB+P treated females only (Tukey post hoc test, p<0.0135, “peak” compared to other time points). Therefore, despite elevated LH basal levels, C451A-ERα mice appear able to mount a LH surge.

Three weeks later, the same mice were then subjected to the same protocol with minor changes. Their blood and brain were collected between 30 min and 1 hour after lights off to evaluate the impact of the mutation on the neuronal circuits underlying the induction of a LH surge by estrogens. As before, C451A-ERα showed higher LH concentrations than WT-ERα (data not shown). The analysis of blood samples collected at euthanasia identified an increase in LH in WT-ERα females treated with EB+P, but not with veh+EB compared to veh+veh (K=10.02, p=0.0067, Fig 2B). By contrast, although LH significantly decreased after EB alone, there was no effect of EB+P in C451A-ERα females (K=7.301, p=0.0260; veh+veh vs EB+veh, p=0.0145; Fig 2B). Comparisons between genotypes in each condition confirmed the higher LH levels measured in C451A-ERα compared to WT-ERα females in all conditions, but not in EB+P condition (veh+veh: U=0, p_adj_<0.0003 EB+veh: U=21, p_adj_=0.0042; EB+P, U=44, p_adj_=0.1137). Contrasting with the observation obtained following repeated blood sampling, these results indicate that only EB+P induces a LH surge in WT-ERα females, but not in C451A-ERα mice.

The brains of these females were then immunostained for Kp and GnRH along with Fos to determine the effect of the mutation on the activation of the hypothalamic circuits underlying the LH surge (49,64). The analysis of the total number of Kp neurons in the AVPv-PeN continum revealed a reduced number of Kp neurons in C451A-ERα females compared to their WT-ERα littermates (F_1,70_ = 38.61; p <0.0001; Fig 2C) and a trend towards an effect of treatment (F_2,70_ = 3.124; p =0.0502). There was however no interaction between the two factors (F_2,70_ = 1.177; p =0.3142). By contrast, GnRH neurons were slightly more abundant in the POA of C451A-ERα compared to WT-ERα mice (F_1,69_ = 5.476; p =0.0222; Fig 2E), but there was no effect of treatment (F_2,69_ = 0.3376; p = 0.7147) or interaction between the two factors (F_2,69_ = 1.976; p = 0.1463).

The analysis of the percentage of Kp and GnRH neurons co-labeled with Fos revealed a very different pattern of response between genotypes. In WT-ERα, EB administered alone or along with P activated a higher percentage of Kp neurons. By contrast, only EB+P elicited such an increased in C451A-ERα females. A 2-way ANOVA indeed identified a trend towards a genotype effect (F_1,70_ = 3.735; p=0.0573), as well as a treatment effect (F_2,70_ = 56.88; p<0.0001) and an interaction between the two factors (F_2, 70_ = 11.17; p <0.0001; Fig. 2D).

Similarly, the percentage of GnRH neurons co-labeled with Fos increased after EB alone and EB+P in WT-ERα females, while only EB+P resulted in such an increased in C451A-ERα females, which resulted in a genotype effect (F_1,69_ = 8.481; p=0.0048), a treatment effect (F_2,69_ = 34.52; p <0.0001) and an interaction between the two factors (F_2,69_ = 3.458; p=0.0371; Fig. 2F). Together, these results indicate that, while EB alone and EB+P activate Kp and GnRH neurons in WT-ERα females, only the EB+P combination mimics these effects in C451A-ERα females.

The percentages of activated Kp and GnRH neurons correlate with circulating LH concentrations in WT-ERα females treated with EB+P (Kp, R^2^= 0.3987; GnRH, R^2^= 0.7037), while it is not the case for the circulating LH in C451A-ERα females (C451A-ERα, Kp, R^2^=0.0157; GnRH; R^2^= 0.0125, data not shown). This difference could be explained by the fact that brains and bloods were collected between 30-60 min of lights off in the second run, while the highest LH concentration measured in the first run were observed at lights off in WT-ERα females but an hour and a half later in C451A-ERα females. Therefore, although EB+P increases the number of Kp and GnRH neurons in C451A-ERα females, this neuronal activity may be associated with a delayed surge of LH.

### Does E_4_ block the LH surge induced by estradiol benzoate (EB)?

Based on the lack of LH surge in C451A-ERα females treated with EB alone but not with EB+P, we postulated that mERα activation is required for the induction of a LH surge by EB but that P can somehow bypass the effect of mERα. As E_4_ was described as an antagonist of mERα (41), we wondered whether E_4_ could block the LH surge induced by EB and whether this effect could be prevented by P.

This experiment followed a similar design as the second part of the previous experiment, except that five treatments (veh+P, EB, EB+P, EB+E_4_, EB+E_4_+P) were compared in wild-type mice (Fig. 3A). As expected, LH levels assayed on samples collected before treatment (day 8) did not differ between groups (F_4,40_=0.4620, p=0.7631, Fig. 3B). By contrast, LH levels assayed within 1 hour of lights off (28 hours after treatment; day 9) were significantly elevated in females treated with EB+P and EB+E_4_+P compared to controls (veh+P), but not in females treated with EB+veh and EB+E_4_ (F_4,39_=11.76, p<0.001; Fig 3C).

The brains of these females were immunostained for Kp or GnRH along with Fos to determine the effect of E_4_ on the activation of the hypothalamic circuits underlying the LH surge (49,64). The total number of Kp neurons in the AVPv-PeN continum (Fig 3D) and preoptic GnRH neurons (Fig 3F) did not differ between treatments (Kp: F_4,31_=0.4461, p=0.7744; GnRH: F_4,33_=0.4645, p=0.7612) but the percentage of Kp and GnRH neurons expressing Fos differed between treatments (Kp, F_4,31_=6.710, p=0.0005; GnRH, F_4,31_=8.489, p<0.0001; Fig 3E-G). These effects resulted from the significantly higher percentage of activated Kp neurons compared to the control condition (veh+P) observed following the administration of all combinations of EB with or without E4/P with the exception of EB combined with only E_4_.

Together, these results indicate that in the absence of exogenous progesterone E_4_ prevents the activation of the neural circuit underlying the induction of an LH surge.

### Chronic E_4_ treatment decreases the number of oocytes ovulated as well as the number of pups per litter

To determine whether a chronic treatment with E_4_ blocked ovulation in mice as reported in rats (45), naturally cycling wild-type females were treated for 3 days with E_4_ (6, 20 and 60µg), E_2_ (0.6µg) as positive control, or veh as a negative, and placed overnight with a male (Fig 4A). The treatment was prolonged for 2 days after mating. This procedure was repeated four times with an interval of 2 weeks so that each female was finally exposed to all treatments in a counterbalanced order, except for females treated with E_2_ since this group was introduced only during the last test. The analysis of the number of females with a vaginal plug revealed that E_2_ and the two highest doses of E_4_ increased the percentage of females with a plug more than 2.5 fold for E_2_ and more than 3.5 fold for E_4_ (60µg) (Fisher’s exact test, veh vs E_2_, p_adj_ =0.0196; veh vs E_4_ (6µg), p_adj_ > 0.9999; veh vs E_4_ (20µg), p_adj_ =0.0004; veh vs E_4_ (60µg), p_adj_ <0.0001; Fig 4B).

**Fig. 4.**
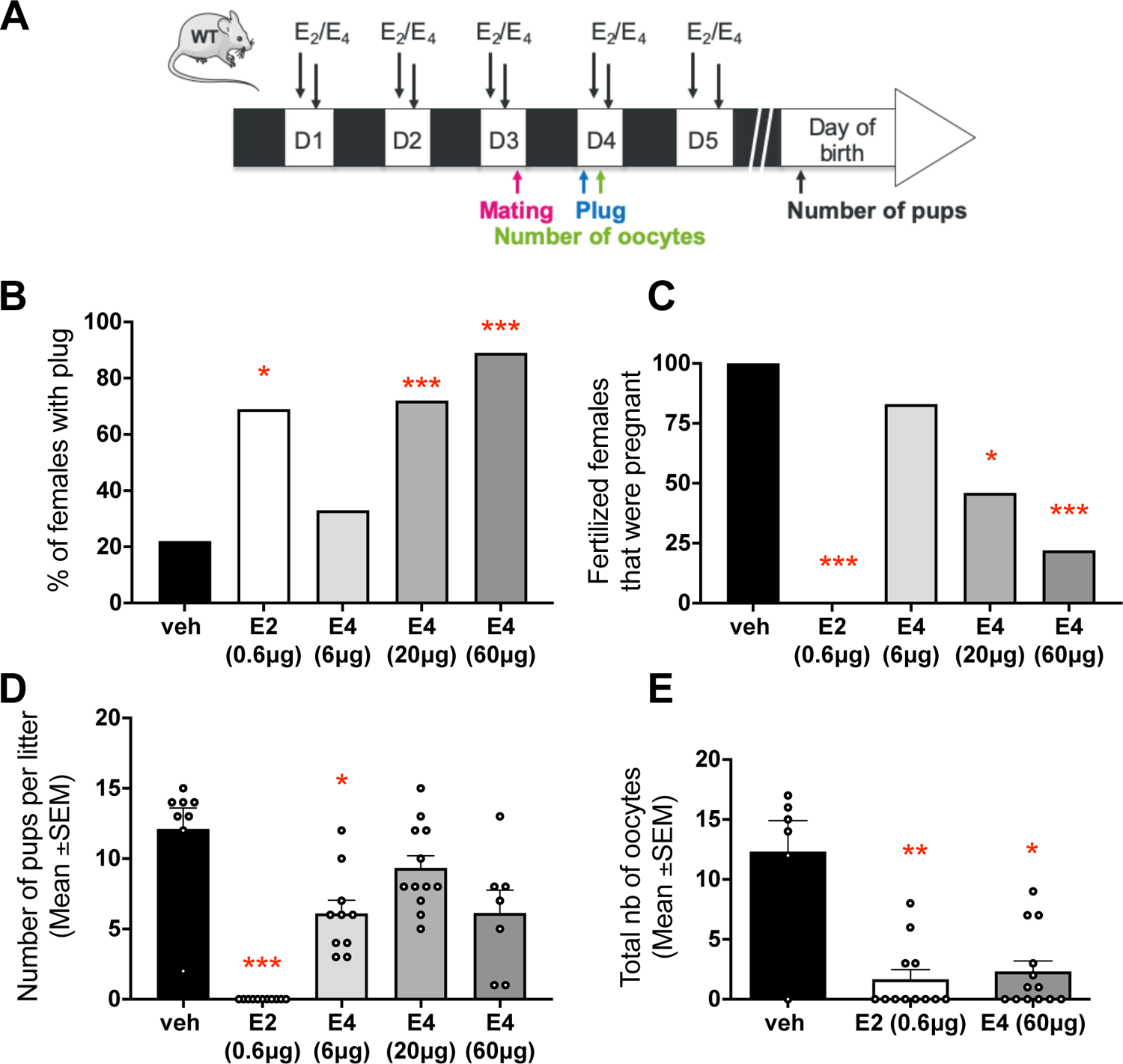
Effect of chronic treatment with estetrol compared to estradiol on the fertility gonadally intact wild-type mice. **A.** WT intact mice were chronically treated with estrogens (black arrows) by subcutaneous injections twice a day for five days. The black arrows represent either an injection of E_2_ (30µg/kg), E_4_ (3mg/kg) or oil and animals were tested either for presence of a plug, pregnancy), the number of birth or for the number of oocytes counted in day 4. **B.** Estetrol (E_4_) dose-dependently increases the percentage of gonadally intact females with a plug after one night of mating (Fisher exact test: *, *** p<0.01, 0.001 vs veh). **C.** E_4_ administered 3 days prior to and 2 days post-mating dose-dependently decreases the percentage of pregnant females (Fisher exact test: *** p<0.001 vs veh). **D.** E_2_ and the lowest dose of E_4_ reduced the number of pups per litter (Kruskal Wallis test; each dot represents one mother). **E.** E_2_ and E_4_ reduce the number of oocytes counted in the two ampulae (Dunn’s test following significant Kruskal Wallis test).

The analysis of the number of mated females that became pregnant and gave birth to pups indicated that E_2_ reduces fertility, as none of the fertilized females became pregnant and delivered pups, while control females all did (Fisher’s exact test: p_adj_ < 0.001; Fig. 4C). Similarly, E_4_ dose-dependently reduced the percentage of fertilized females that became pregnant and gave birth (veh vs E_2_ (0.6µg), p_adj_<0.001, veh vs E_4_ (6µg), p_adj_>0.9999; veh vs E_4_ (20µg), p_adj_=0.0204; veh vs E_4_ (60µg), p_adj_<0.001).

Interestingly, treatment also affected the number of pups per litter (K=32.97, p<0.001), which was abrogated by E_2_ (p<0.001) and also decreased by the lowest dose of E_4_ compared to vehicle (p=0.0495; Fig 4D). Together, these observations indicate that E_4_ could impair ovulation as supported by previous work in rat (45). However, it is also possible that E_4_ impairs embryonic development as the absence of membrane signaling of ERα was recently shown to interfere with placentation leading to infertility (39).

To discriminate between these two possibilities, the same females were then divided in three groups treated for 3 days with E_2_ (0.6µg), the highest dose of E_4_ previously tested (60µg) or vehicle. On the morning following mating with a male, the number of cumulus oophori ovulated were counted in females displaying a plug. Although less females treated with E_2_ (33%) and E_4_ (54%) ovulated compared to control females (80%), these proportions did not differ significantly, probably due to the low sample size (Fisher’s exact test, veh vs E_2_, p=0.1312; veh vs E_4_, p=0.3331). By contrast, E_2_ and E_4_ significantly reduced the number of oocytes counted in the two uterine horns (K=9.576, p=0.0083; veh vs E_2_, p=0.0076; veh vs E_4_, p=0.0332; Fig 4E). These data indicate that E_2_ or E_4_ provided around mating inhibit ovulation, thus adding to the contraceptive effect of E_4_ reported in rats and humans (45,46).

### E_4_ mimics the effect of E_2_ for lordosis but not for partner preference

The observation that gonadally intact females treated with E_4_ are more likely to present a plug after an overnight interaction with a male (Exp.3; Fig. 4B) suggested that E_4_ stimulates sexual proceptivity and/or receptivity. To address this question, OVX females were treated twice a day with E_2_ (0.5µg) and/or E_4_ (60µg) for two days and received the estrogen again plus progesterone 3-4 hours prior to being tested for lordosis or partner preference (Exp.5; Fig 5A). This dose of E_2_ combined with P is known to induce lordosis behavior in OVX females, while the dose of E_4_ can antagonize mERα while mimicking the nuclear actions of E_2_.

**Fig. 5.**
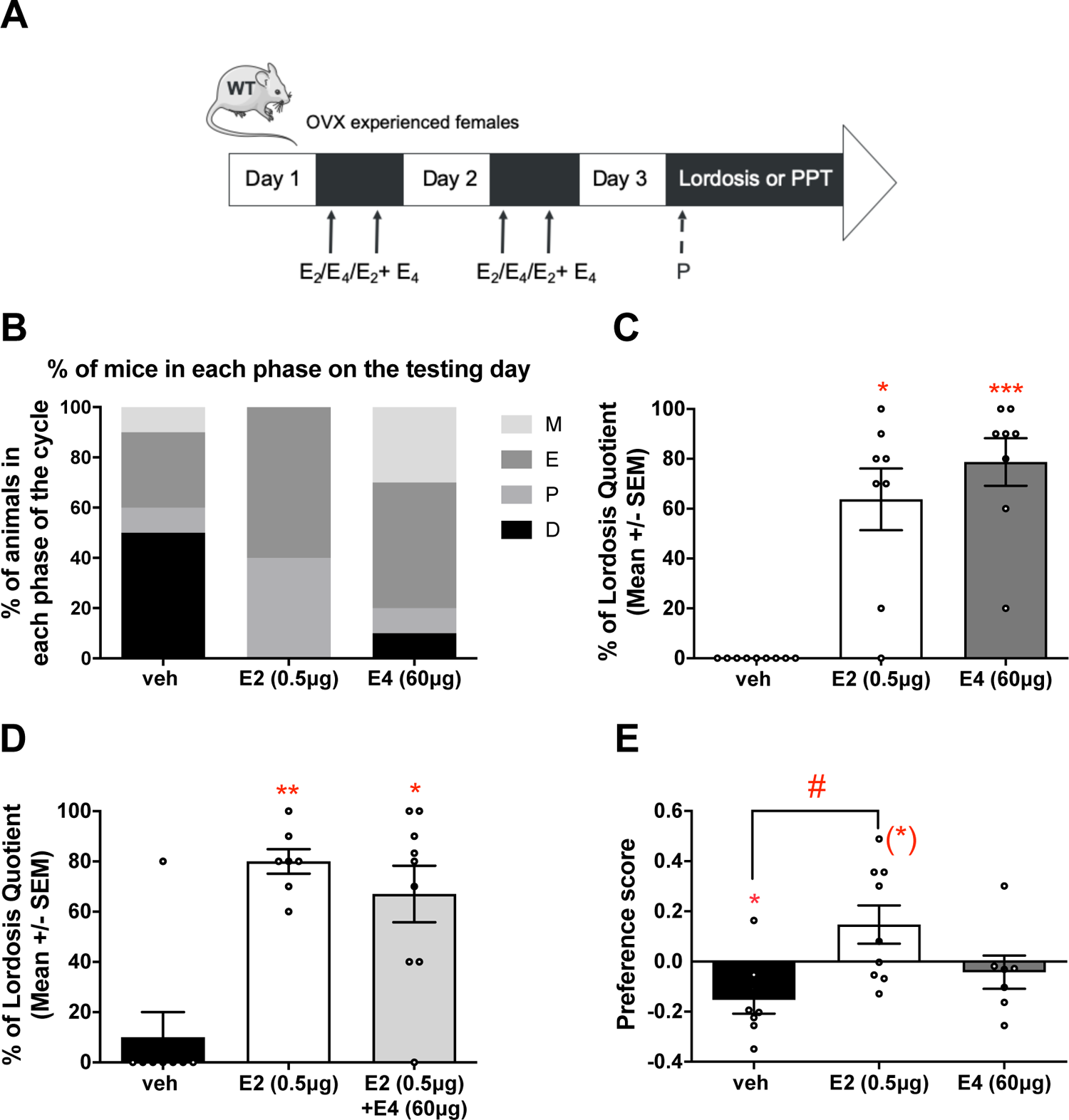
Effect of chronic treatment with estetrol compared to estradiol on sexual behavior of gonadally intact wild-type mice. **A.** OVX experienced females were chronically treated with estrogens (black arrows) by subcutaneous injections twice a day for two days. The third day all mice received an injection of progesterone (P (25mg/kg); dashed arrow) excepted when tested for cyclicity and were tested for either cyclicity (smeared every day), lordosis or partner preference (PPT). **B.** E_2_, but not E_4_, alters the proportion of females in each phase of the cycle after 3 days of treatment (chi square; M: metestrus, E: estrus, P: proestrus, D: diestrus; E and P as well as M and D were pooled for the analysis). **C.** Both E_2_ and E_4_ increased lordosis quotient (Kruskal Wallis test: *,*** p<0.05, 0.001). **D.** E_4_+E_2_ induce the same effect as E_2_ alone (Kruskal Wallis test: *,** p<0.05, 0.01). **E.** While OVX females preferred spending time with a female (negative score), females treated with E_2_ tended to prefer the male (positive score). However, females treated with E_4_ did not show a preference (one sample t test, (*), * p<0.10, 0.05 vs 0; Kruskal Wallis test: # p<0.05).

Vaginal smears were carried out to evaluate the impact of the treatment on the estrous cycle. The proportion of females in proestrus or estrus significantly differed between treatments (X_2_ = 8.4, p = 0.015; Fig 5B). Fisher exact tests revealed that E_2_ increased the proportion of females in proestrus or estrus compared to veh (p = 0.011), while it was not the case for E_4_ (p = 0.656).

A first test revealed that when given separately both E_2_ and E_4_ significantly induce lordosis behavior compared to OVX controls (K = 16.59, p = 0.0002; Dunn’s post-hoc test vs veh: E_2_, p = 0.01; E_4_, p = 0.0003; Fig 5C). A second test evaluated whether E_4_ could antagonize the effect of E_2_ and showed that E_2_+E_4_ induces the same effect as E_2_ alone (K = 10.97, p = 0.004; Dunn’s post-hoc test vs veh: E_2_, p = 0.0099; E_2_+E_4_, p = 0.016; Fig 5D), thus suggesting that E_4_ acts here as an ER agonist.

Finally, a last test compared the effect of E_2_ and E_4_ alone on partner preference. Non-directional one-sample two-tailed t tests revealed that vehicle treated females spent more time with the stimulus female (t_7_ = 2.772, p = 0.028; negative score = preference for the stimulus female), while E_2_ treated females tended to prefer spending time with the male stimulus (t_8_ = 1.938, p = 0.089). However, E_4_ treated females did not show any preference for one sex (t_6_ = 0.645, p = 0.543; Fig 5E). These results were confirmed by a Kruskal-Wallis test (K = 8.138, p = 0.017) revealing a significant difference between females treated vehicle and E_2_ (p = 0.033) but not with E_4_ (p > 0.432).

## Discussion

The goal of the present study was to investigate the contribution of membrane estrogen receptor alpha (mERα) to the positive feedback of estrogens on LH secretion. We found that both a mouse model constitutively lacking mERα signaling (38) and a natural estrogen described to have an antagonistic action on mERα (41) blocked the ability of E_2_ to induce a LH surge and the neural circuit associated to this physiological response in the absence of progesterone, thus suggesting that mERα plays a role in this process. Moreover, we confirmed that, in mice like in rats, E_4_ dose-dependently blocks ovulation. Along with its action on the positive feedback, this observation provides additional evidence of the antagonistic action of E_4_ on membrane ERα in the brain. Finally, we found that E_4_ also mimics the stimulatory effect of E_2_ on lordosis behavior.

### C451A-ERα females show a distinct phenotype of LH secretion

The idea that the positive feedback of estrogens depends on nuclear estrogen signaling is mainly based on the observation that the induction of a LH surge requires a prolonged exposure to high estrogen levels (24,25). Moreover, restoring ERE-independent ERα signaling had failed to restore the capacity to mount a LH surge in response to estrogens in ERαKO mice indicating that non-classical signaling alone is not sufficient for positive feedback (26). However, previous evidence support the existence of a cooperation between nuclear- and membrane-initiated estrogen signaling (65,66). Therefore, it is likely that membrane estrogen signaling requires nuclear estrogen signaling to properly function even if classical signaling constitutes the prime requisite. To test this possibility, we used two complementary approaches. With a point mutation at the site of palmitoylation of ERα, C451A-ERα mice allow the study of the impact of a lack of membrane signaling of ERα while preserving its nuclear activity (38). Although this mutation does not seem to alter the sexual differentiation of females (67), the constitutive absence of mERα is an obvious limitation of such a model that may result in developmental defects as well as compensations. On the other hand, the antagonistic action of E_4_ on the membrane estrogen signaling of the classical estrogen receptors ERα (and possibly ERβ) provides a means to circumvent developmental deficits or compensation. E_4_ has obviously mixed actions on the nuclear and membrane signaling of ERs limiting our possibility to draw definitive conclusions. Moreover, its preference for ERα over ERβ as well as its properties in the brain remains poorly documented. However, the comparison of effects obtained with both approaches provides confidence that membrane signaling plays a role in the positive feedback of estrogens on LH secretion.

Gonadally intact C451A-ERα females exhibit elevated LH levels and fewer corpus lutea than wild-type females (38), suggesting a potential role of mERα signaling in both negative and positive feedbacks. The C451A mutation does not alter brain expression of ERα and Kp (67) supporting the preserved transcriptional activity of the nuclear fraction of the receptor. The present study reveals however that C451A-ERα mice exhibit a distinct profile of LH secretion in response to estrogens compared to other ERαKO mouse models. Indeed, contrasting with ubiquitous ERαKO, neuron-specific ERαKO and ARC specific ERαKO mice, which exhibit altered LH responses to ovariectomy and/or E_2_ (6,68,69), C451A-ERα females respond to both ovariectomy and provision of exogenous estrogens (Fig 1B; Faure et al., in preparation). However, they seem to be less sensitive to circulating E_2_ levels since they require higher doses of E_2_ to restore basal LH blood concentrations (Faure et al., in preparation). C451A-ERα females were also unable to respond to increasing E_2_ levels by a LH surge (Fig. 1 and Fig. 2). This is congruent with previous observations in other ERαKO or knock down models (7,22,70–72). Interestingly, when treated with P, C451A-ERα females exhibit surge-like changes in LH levels measured in repeated blood sampling. Similarly, despite an apparent absence of change in LH levels compared to vehicle treated controls, they display the typical activation of Kp and GnRH neurons following a positive feedback induction protocol including P (49,50). Although one might wonder how a LH surge is possible in mice showing such elevated circulating LH concentrations, this idea is compatible with the recent “two component model” of control of GnRH secretion which poses that the positive and negative feedbacks of gonadal steroids on LH regulation are regulated by two anatomically distinct and independent mechanisms (Herbison 2020). This idea is also supported by a recent study showing that a surge can be elicited over high LH levels (73).

The prevention of EB-induced rise in LH and the activation of Kp and GnRH neurons by E_4_ (Fig. 3) confirms the effect of a lack of mERα. The timing of these effects does not allow to determine whether they reflect direct membrane actions or membrane-initiated transcriptional effects. Further studies will be needed to identify the mechanism underlying these effects.

### E_4_ acts as a mERα antagonist in the brain

E_4_ mimics the nuclear actions mediated by E_2_ on ERα in several tissues (41-43,74). Although antagonistic actions of E_4_ have been reported in several tissues including the brain, whether they rely on transcriptional or membrane actions is not known with the exception the membrane-mediated action identified in endothelial cells and in breast cancer cells (41,44,74–77). The blockade of E_2_-induced LH surge by E_4_ in parallel with the absence of LH surge in mice lacking mERα signaling provide strong evidence of the antagonist action of E_4_ on mERα in the brain, and most probably within the preoptic area.

### Discrepancies between the two approaches

Although the two approaches employed in this study lead to similar conclusions, differences were observed. E_4_ altered EB-induced activation of Kp and GnRH neurons and LH surge (Fig. 3) but had no impact on the number of Kp and GnRH neurons. By contrast, OVX and E_2_-treated C451A-ERα females exhibited elevated LH concentrations along with fewer Kp neurons and more GnRH neurons than their wild-type counterparts (Fig. 2). The presence of Kp in AVPv neurons of C451A-ERα mice confirms the preserved transcriptional activity of ERα, since a complete absence of ERα in Kp neurons leads to an absent or greatly reduced Kp expression (70,78,79). However, the lower number of Kp neurons observed in the present experiment could be explained by a developmental effect of the constitutive mERα absence or by an effect of the mutation on Kp transcription. Although developmental defects cannot be ruled out, several observations points to the latter. First, the early programming of AVPv Kp neurons is affected by estrogen exposure in two critical periods. Perinatal exposure to estrogens leads to few detectable Kp neurons that are typical of males (49). In females, prepubertal exposure to estrogens is required to observe normal adult Kp neuronal numbers (50,80,81). A previous study showed that C451A-ERα females, but not males, exhibit expected numbers of Kp in the AVPv supporting an absence of programming defect in this cell population in females (67). Second, the number of Kp neurons in the AVPv of C451A-ERα females appears to be influenced by the dose of estrogens. Comparable numbers of Kp neurons were counted in the AVPv of wild-type and C451A-ERα females injected daily with EB (1µg) for 2 weeks (67). Moreover, a parallel study found a difference between genotypes in females implanted with a Silastic capsule filled with 1µg of E_2_ but not with capsules containing 5µg of E_2_ (Faure et al., in preparation). ERα may thus be less sensitive to estrogens in the mutant mice thus requiring higher circulating concentrations of E_2_ to stimulate normal Kp expression (Faure et al., in preparation). Finally, ERE-independent pathways are not sufficient to stimulate Kp expression in the AVPv of ERαKO mice (78). The lower number of Kp neurons in C451A-ERα mice thus seems attributable to a lower expression of Kp in the presence of low circulating estrogens. One report mentions, however, a stimulatory role for mERα in the expression of Kp in mHypo51A cells, an immortalized line derived from adult mouse hypothalamic neurons presumed to be AVPv Kp neurons (37). The lack of effect of E_4_ of on the number of AVPv Kp neurons argues against this conclusion.

### Role of progesterone signaling

Genetic or pharmacological blockade of mER signaling prevented the induction of a LH surge by EB. In both cases, LH surge was restored by the administration of P 3 to 4h before lights off. The potentiating effect of P on EB-induced surge has long been known (62). The importance of P for the induction of the LH surge is underlined by studies focusing on progesterone receptors (PR), whose expression is stimulated by estrogens through an ERE-dependent genomic action (82). Knockout PR mice (PRKO) and mice lacking PR exclusively in Kp neurons (KissPRKO) are unable to mount an EB-induced surge (83–85). However, the reintroduction of PR expression specifically in Kp neurons of the AVPv of KissPRKO mice recovers the LH surge, demonstrating the essential role of P action on this neuronal population for the induction of the LH surge (86). Our results could thus suggest that the absence or blockade of mERα impedes PR expression. This hypothesis seems however unlikely given that C451A-ERα mice respond well to exogenous P in terms of Kp and GnRH activation. Moreover, E_4_ mimics the action of E_2_ on PR expression and E_2_ induces PR expression in the brains of C451A-ERα females, albeit to a lesser extent than in wild-type mice (Faure et al., in preparation). Membrane estrogen signaling could also interfere with another aspect of P signaling, such as its membrane-initiated or ligand independent signaling (87). Alternatively, mERα could modulate local P synthesis to contribute to LH surge induction. Remarkably, all the studies supporting the necessity of PR, in particular within Kp neurons, to induce an LH surge did not provide exogenous P, suggesting that an endogenous source of P exists in OVX females which may be necessary to elicit the surge. This idea is supported by an absence of correlation between circadian fluctuations of brain and plasma P concentration in ovary intact mice (88) and the work of Micevych indicating that a rise in neuroprogesterone produced by hypothalamic astrocytes is a prerequisite for the LH surge (37,89–91).

### Effect of E_4_ on sexual behavior

The increased number of mated females following E_4_ treatment prompted us to investigate the effect of E_4_ on sexual behavior. Females treated with E_4_ were not more attracted to males but showed increased lordosis behavior, while E_2_ stimulated both responses. This discrepancy in the effects of the two estrogens in these two tests may be explained by a different contribution of membrane and nuclear estrogen signaling to the regulation of the two aspects of behavior (motivation vs copulation) as was previously suggested in males ((92); but see also below). It could also be argued that the partner preference test may not reflect the complexity of the motivational aspects and that another measure of sexual motivation that was not tested here may have been affected. Nonetheless, the effect of E_4_ on lordosis behavior suggests that E_4_ enhances female sexual receptiveness, thus uncoupling the stimulatory actions of E_2_ on the preovulatory LH surge and sexual behavior which is critical for reproduction.

The absence of inhibitory action of E_4_ on E_2_-induced lordosis behavior and the normal sexual behavior of C451A-ERα females (67) is however somewhat surprising given previous evidence supporting the need of mERα for proper lordosis behavior in rats (93–95). Our observations may translate a species difference in the role of mERα in this behavior with possibly the involvement of another receptor, like GPER1 (96). It is also possible that the conditions in which our mice were tested do not allow to see the impact of mERα blockade/deficit because exactly like the effect on the LH surge this inhibitory effect is prevented by P, as was previously suggested (67).

## Conclusions

The present results contradict the long-standing idea that the induction of a LH surge by rising concentrations of circulating estrogens is mediated by genomic effects only. Although it has long been known that the LH surge requires a prolonged exposure to high estrogen concentrations, it is also recognized that estrogens do not have to be present the whole time for the surge to occur (24,25). Moreover, membrane estrogen signaling through modulation of intracellular signaling cascades can potentiate the slower transcriptional actions of estrogens (66,97). A role for membrane initiated signaling in the induction of LH surge by estrogens is supported by the acute actions of E_2_ reported on the activity of GnRH neurons (32,35,36). It should also be pointed out that membrane initiated signaling does not necessarily imply rapid actions, as indirect genomic signaling is also possible. The present results cannot discriminate between these possibilities, nor can they determine the contribution of mERα located in the AVPv and ARC. Although it cannot be ruled out that the impaired positive feedback observed in mutant mice is an indirect result of the expected dysregulation of the negative feedback, this would not explain why Kp and GnRH neurons are still able to respond normally when provided with P along with EB. Moreover, the similarity of the responses of C451A-ERα mice to wild-type females treated with E_4_ supports a role for membrane-initiated estrogen signaling in the EB-induced LH surge. Together, the present work provides evidence for a role of membrane estrogen signaling in the induction of the LH surge. Further work will be necessary to identify where this contribution occur. Based on the absence of ERα in GnRH neurons and the prominent role of Kp neurons of the AVPv as drivers of the GnRH surge, this neuronal population constitutes a good candidate.

## Acknowledgements

This work was supported by grants from the Fonds National pour la Recherche Scientifique (F.R.S.-FNRS PDR T.0042.15) and the Special Funds for Research from the University of Liège (<u>FSR-S-SS-19/40</u>) and a research project (E4Liberty) with Mithra Pharmaceuticals. CdB was a Post-doctoral Researcher of the F.R.S.-FNRS and CAC is a Research Director of the F.R.S.-FNRS. We thank Laura Vandries and Céline Roomans for their help with the immunostaining and Arlette Gérard for carrying out the RIA assay.

